# Modeling of HIV-1 prophylactic efficacy and toxicity with Islatravir shows non-superiority for oral dosing, but promise as a subdermal implant

**DOI:** 10.1101/2024.04.15.589489

**Authors:** Hee-yeong Kim, Lanxin Zhang, Craig W Hendrix, Jessica E Haberer, Max von Kleist

## Abstract

HIV prevention with pre-exposure prophylaxis (PrEP) constitutes a major pillar in fighting the ongoing epidemic. While daily oral PrEP adherence may be challenging, long-acting (LA-)PrEP in oral or implant formulations could overcome frequent dosing with convenient administration. The novel drug islatravir (ISL) may be suitable for LA-PrEP, but high doses have been associated with lymphopenia.

We developed a mathematical model to predict ISL pro-drug levels in plasma and active intracellular ISL-triphosphate concentrations after oral vs. subdermal implant dosing. Using phase-II trial data, we simulated antiviral effects and estimated HIV risk reduction for multiple dosages and dosing frequencies. We then established non-toxic exposure thresholds and evaluated low-dose regimens. Our findings suggest that implants with 56–62 mg ISL offer safe, effective HIV risk reduction without being toxic. Oral 0.1 mg daily, 3–5 mg weekly, and 10 mg bi-weekly ISL provide comparable efficacy, but weekly and bi-weekly doses resulted in residual toxicity, while adherence requirements for daily dosing were similar to established oral PrEP regimen. Oral 0.5–1 mg on-demand provided *>* 90% protection, while not being suitable for post-exposure prophylaxis.

## Introduction

Over 40 years after the discovery of HIV as cause of AIDS [1, 2], the epidemic remains a significant public health concern. By 2022, HIV caused over 85 million infections and 40 million deaths [3], one every minute [4]. While highly active antiretroviral treatment can prevent AIDS, there is still no cure for HIV nor an effective vaccine.

Pre-exposure prophylaxis (PrEP) has proven effective in preventing HIV transmission when taken as directed [5]. Currently, three options are available: daily or on-demand oral PrEP with emtricitabine/tenofovir disoproxil fumarate (FTC/TDF) or FTC/tenofovir alafenamide (TAF), injectable long-acting cabotegravir (CAB-LA) and a monthly vaginal ring containing dapivirine (DPV-VR) [6]. Oral FTC/TDF PrEP is widely available in high-income settings, particularly among men-who-have-sex-with-men (MSM) [7], with potential for an about 95% reduction in HIV acquisition risk, even when taken on-demand [8, 9]. In heterosexual women, the success of FTC/TDF PrEP may be hampered by poor adherence [10], and on-demand use is currently not recommended by the WHO [11]. However, CAB-LA PrEP demonstrated an approximately 94% incidence reduction in heterosexual women [12], suggesting long-acting (LA-)PrEP as excellent alternative for those unable to adhere to a daily regimen. The investigational drug islatravir (ISL, MK-8591 or EFdA) may have considerable potential for LA-PrEP, due to its antiviral potency (sub-nanomolar range), long intracellular half-life and high barrier to drug resistance [13, 14]. So far, the pharmacokinetics and antiviral potency of ISL have been assessed for daily, weekly and monthly oral dosing, and as a subdermal implant, which may need to be replaced every 3–6 months.

ISL is a deoxyadenosine analogue and inhibits HIV-1 reverse transcriptase (RT). Unlike approved nucleoside-RT-inhibitors (NRTIs) like FTC/TDF and FTC/TAF, which act as immediate polymerase chain terminators [15, 16], ISL is a first-in-class nucleoside-RT-translocation-inhibitor (NRTTI) [17, 18]. After uptake by HIV target cells and tri-phosphorylation into its active form, islatravir-triphosphate (ISL-TP) [19, 20] competes with natural dATPs during reverse transcription. With a 2-fluoro group, ISL has increased intracellular stability, lasting approximately 78.5–128 hours. It exerts multiple mechanisms of action (MOA): The 4’-ethynyl group acts as a weak chain terminator, blocking further nucleotide incorporation into the nascent pro-viral DNA. If RT translocation still occurs, the 3’-OH group causes conformational distortions of the RT-primer/template complex leading to delayed chain termination after incorporation of 1–3 additional nucleotides [21–23].

In 2021, the U.S. Food and Drug Administration (FDA) halted clinical trials [24] after CD4^+^ T-cells declines were observed in participants receiving ISL in combination with other antivirals [25, 26]. Those having taken ISL alone for PrEP experienced a dose-dependent drop in total lymphocyte counts (lymphopenia) [27]. However, new trials were announced that assess lower doses ISL for HIV treatment [25], including weekly oral administration with doravirine [26], while PrEP trials were discontinued.

We developed an integrated mathematical model of ISL’s pharmacokinetics (PK), pharmacodynamics (PD), prophylactic efficacy, and toxicity using phase I–II clinical data. This model can help guide further consideration of ISL for HIV treatment and/or prevention. We simulated multiple dosing schemes for both oral and implant administration, determined dose-dependent adverse effects, and assessed toxicological risks and prophylactic efficacy of low-dose ISL regimens. Our approach may identify a safe and effective niche for ISL, serving as a blueprint to guide the further development of NRTTIs for HIV prophylaxis and treatment.

## Methods

### Pharmacokinetic (PK) data

We considered all relevant, publicly available (pre-)clinical data on ISL, including oral and implant formulations [19, 20, 28–30]. For oral doses of 0.5–400 mg, this included 10 datasets on average plasma ISL PK and 11 datasets on intracellular ISL-TP in peripheral blood mononuclear cells (PBMCs), which was used as marker for the antiviral effect-site in HIV PrEP. For subdermal implant administration, we examined 5 loading doses with ISL levels ranging from 48–62 mg, including two doses (54 and 62 mg) reporting both plasma ISL and intracellular ISL-TP [20, 29].

### PK modeling

We considered a semi-mechanistic compartment model to predict plasma ISL and intracellular ISL-TP pharmacokinetics as depicted in Fig. 1A with ordinary differential equations outlined below.

**Fig 1.**
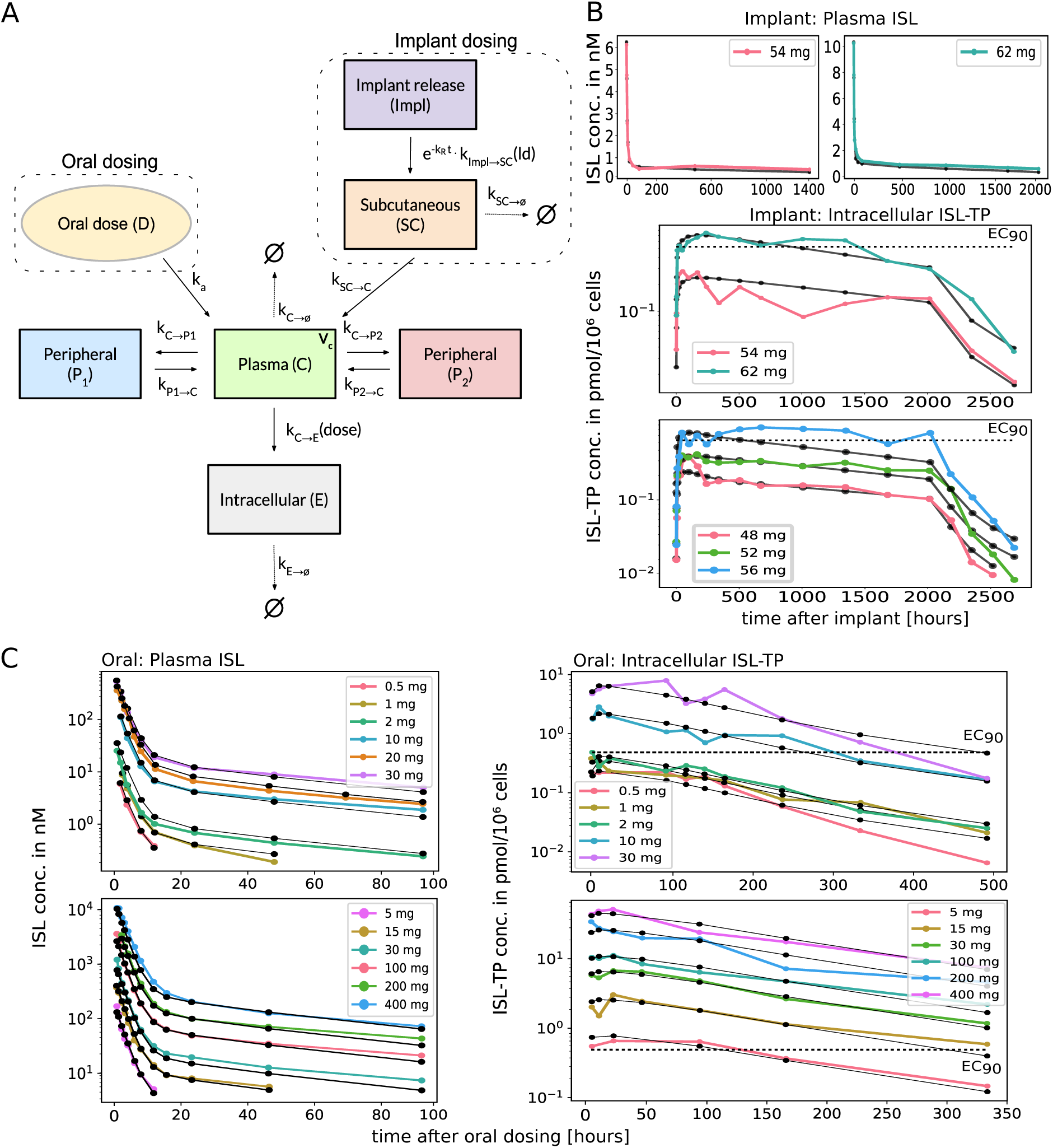
Pharmacokinetics of ISL. A. Integrated pharmacokinetic model encompassing the two routes of drug administration (oral and implant). B. Clinical (colored-dotted lines) and corresponding model-predicted (black-dotted lines) plasma ISL and intracellular ISL-TP concentrations in PBMCs for implants with different loading doses. Implants were removed after 12 weeks. Data was derived from [20, 29]. C. Clinical (colored-dotted lines) and corresponding model-predicted (black-dotted lines) plasma ISL and intracellular ISL-TP concentrations after single oral doses. Data was derived from [19, 28, 30]. For reference, model-computed 90% HIV-protective intracellular ISL-TP concentrations (EC_90_) are shown as dashed horizontal lines.

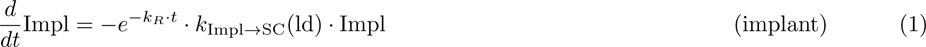

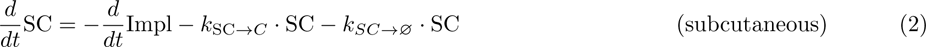

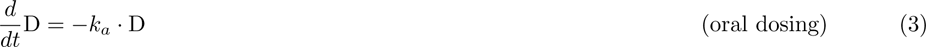

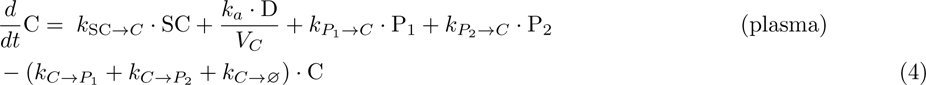

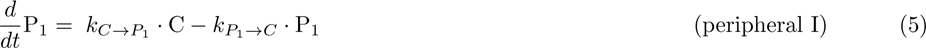

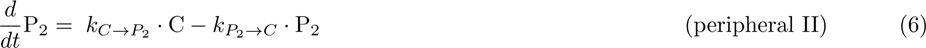

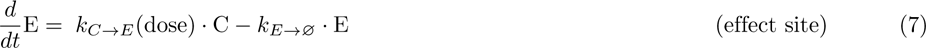

### Implant administration

Subdermal ISL implants are placed on the inner side of the non-dominant upper arm. In our PK model, they release ISL (Impl) into the central (plasma) compartment ‘C’ via the subcutaneous compartment (SC), Fig. 1A. The cumulative drug release from the implant, estimated from an *in vitro* experiment [31] for non-erodible EVA (ethylene vinyl acetate) at 54 mg and 62 mg loading doses, follows a simple exponential function 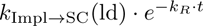, where ‘ld’ denotes the loading dose. Rates for 48 mg, 52 mg, and 56 mg were linearly interpolated in line with [31] (Supplementary Table S1). Concentration transfer SC *→* C was modeled by first-order kinetics with rate *k*_SC_*_→C_*. Additionally, a perceptible clearance in the subcutaneous compartment at rate *k_SC→_*_∅_ was included based on the data.

### Oral drug dosing

We modeled first-order absorption kinetics with rate parameter *k_a_*from the oral dosing compartment D into the central plasma compartment C, in line with previous studies [18].

### Plasma pharmacokinetics

We considered three compartments (central plasma C, and two peripherals P_1_, P_2_) to model ISL plasma PK after oral or implant dosing. The ISL plasma PK shows a tri-phasic profile, with two phases evident for oral data (Fig. 1C) and the third phase visible only after implant administration (Fig. 1B). For the central compartment, we set its volume to the physiological volume of blood plasma (*V_C_* = 3.5 L) [32]. The rate parameters for drug transfer between C and P_1_ are denoted by 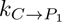 and 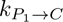, respectively. Analogously, 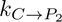 describes the rate constant of ISL transfer of C to P_2_, and 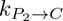 the transfer back. The elimination rate constant in the central compartment is denoted by *k_C→_*_∅_.

### Intracellular pharmacokinetics

The effect compartment E reflects the concentration of ISL-TP in PBMCs. Influx of plasma ISL into target cells and intracellular conversion to ISL-TP is modeled as dose-dependent input rate parameter *k_C→E_*(dose), while elimination of intracellular ISL-TP is modeled by first-order kinetics with parameter *k_E→_*_∅_. We ignored any flux of ISL from the cellular into the central compartment, which would only marginally affect plasma PK, due to the very small volume of the PBMC compartment (*≈* 10*^−^*^6^ L) [33].

### Initial Conditions

Oral dosing was simulated with the respective initial doses in the dosing compartment D, while all other compartments were either zero (single dose) or set to the pre-dosing concentrations (multiple dose). For implant administration, an initial fraction *π*_ld_ of the loaded drug ld *∈ {*48, 52, 54, 56, 62*}* mg was assigned to the subcutaneous tissue, while the rest (1 *− π*_ld_) remained in the implant. This drug fraction corresponds to small damage to both implant and tissue, upon insertion. Implant data was available for first-(54, 62 mg) and second-generation (48, 52, 56 mg) formulations. Parameters *π*_54_ and *π*_62_ were estimated from first-generation data, comprising plasma and drug release information. The study indicated that the 56 mg implant aimed for similar release rates as the prototype for 62 mg [29]. Consequently, we grouped the remaining *π*_ld_ values into drug loads *≤* 54 mg and *>* 54 mg (Supplementary Table S1).

### Viral dynamics and PK-PD link

To infer antiviral effects of ISL, we used an established HIV-1 viral dynamics model [32]. In essence, this model comprises free infectious virus *V*, as well as early and late infected T-cells and macrophages *T*_1_*, T*_2_*, M*_1_ and *M*_2_. In the viral kinetics model, the process of reverse transcription (hence the transition *V → T*_1_) is inhibited via an Emax-model [34]

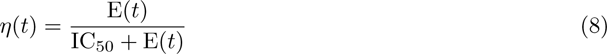

where *η*(*t*) denotes the time-dependent strength of inhibition and E(*t*) denotes the drug concentration at the effect site, compare Eq. (7).

### Prophylactic efficacy

After estimating viral load (VL) kinetics data based on the HIV-1 viral dynamics system, a reduced model that is sufficient to accurately predict prophylactic efficacy *φ* [35] was used. *φ* describes the reduction in infection risk per contact, which can be estimated after coupling ISL PK to the viral dynamics model [32].

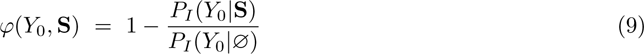

Infection probabilities can be computed for a particular drug regimen **S** or without any drug, denoted as *P_I_* (*Y*_0_*|S*) and *P_I_*(*Y*_0_*|*∅), respectively. The initial viral state *Y*_0_ = [*V, T*_1_*, T*_2_] consists of the number of viruses *V*, early infected T-cells *T*_1_ and productively infected T-cells *T*_2_ [32]. We used a recently developed numerical method [35] to estimate *P_I_* (*Y*_0_*|S*) and hence the efficacy of PrEP/PEP for any **S** within seconds.

### PK parameter estimation

In a two-step procedure, we estimated all PK parameters (Eq. (1)–(7)) based on concentration-time profiles shown in Fig. 1B–C using the *trust region reflective method* to minimize the *squared error* between data and model-prediction.

Parameter boundaries were determined through random parameter seeding and multiple local optimizations to obtain parameter ranges for further optimization. We used parametric bootstrap to obtain estimates of parameter uncertainty: Samples were drawn for each point in a dataset of size *n*, assuming a normal distribution with 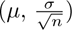. The generated samples were passed to the minimizer until convergence. Resulting parameter estimates are depicted in Table 1. Parameter fitting used plasma drug concentration for implants without transformation, while the remaining data was fitted after log-transformation (Fig. 1B–C). To facilitate parameter uncertainty quantification by parametric bootstrap, we extracted error estimates in the data and assumed that the data was normal distributed 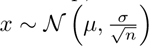, where 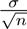 denotes the sampling error of the data and *n* denotes the number of samples contributing to the concentration measurement. To address missing values, interpolation was conducted using the coefficient of variation 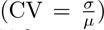 within or between datasets. To estimate bootstrap confidence intervals, we generated 1000 bootstrap samples for each data point and re-fitted the model to each re-sampled dataset. Implant-specific parameters *k_R_, π*(ld)*, k*_Impl_*_→_*_SC_(ld) were not included in the procedure. We validated our model using oral ISL data of 0.25–5 mg once-daily [36] and 60, 120 mg once-monthly [37], focusing on single-dose experiments and implants, Supplementary Figure S1.

**Table 1.** Pharmacokinetic parameter estimates (mean *±* standard deviation). Rate constants are given in the units *h^−^*^1^, the plasma volume *V_C_* was fixed to 3.5L. The first three parameters depend on the route of administration as noted, the remaining parameters are independent of dosing route. A detailed description of the rate constants can be found in the *Methods*.

| Parameter | Value | Parameter | Value |
| --- | --- | --- | --- |
| $k_a$ | $0.4232 \pm 8.2 \cdot 10^{-3}$ (oral) | $k_{C \rightarrow P_1}$ | $1.8494 \pm 9.0 \cdot 10^{-2}$ |
| $k_{SC \rightarrow C}$ | $0.0151 \pm 9.0 \cdot 10^{-4}$ (implant) | $k_{P_1 \rightarrow C}$ | $0.0023 \pm 1.0 \cdot 10^{-2}$ |
| $k_{SC \rightarrow \emptyset}$ | $0.1521 \pm 1.1 \cdot 10^{-2}$ (implant) | $k_{C \rightarrow P_2}$ | $5.2726 \pm 1.7 \cdot 10^{-1}$ |
| $k_{C \rightarrow \emptyset}$ | $6.8999 \pm 1.9 \cdot 10^{-1}$ | $k_{P_2 \rightarrow C}$ | $0.0237 \pm 1.4 \cdot 10^{-3}$ |
| $k_{E \rightarrow \emptyset}$ | $0.0088 \pm 3.0 \cdot 10^{-4}$ | $V_C$ | 3.5 L (fixed) |

### PD parameter estimation

To assess ISL’s potency, we digitized VL data from phase II 10-day monotherapy trial in HIV-infected subjects (*N* = 30) [19, 28]. Using an HIV-1 viral dynamics model (specifications see [32]), we initially computed steady-state levels in the absence of ISL (pre-treatment condition, baseline). By coupling ISL PK with the virus dynamics model (PD) using Eq. (8), we estimated the log-change in VL from (pre-treatment) baseline by fitting the drug potency IC_50_ to the data [19,28]. The fitting process employed the *trust region reflective method*.

### Toxicity thresholds

ISL has been associated with dose-dependent lymphopenia [38] and no mitochondrial toxicity as seen in some NRTIs. The underlying mechanisms are unclear.

In our modeling, we assume that toxic effects correlate with plasma ISL levels (details in *Discussion*). Daily oral dosing of 0.25 mg ISL showed no changes in lymphocyte counts [38]. We reproduced the experiment from that study using our PK model and estimated the maximum plasma concentration, serving as no-observed-adverse-effect-level (NOAEL). With this threshold, we calculated the percentage of time above NOAEL for various dosing regimens.

### Code availability

Codes were written in Python, version 3.10.12, utilizing the scipy.optimize 1.11.1 and scikit-learn 1.3.0 package and are publicly available at https://github.com/KleistLab/ISL_PK-PD-PrEP.

## Results

### PK model building and parameter estimation

Fig. 1A illustrates the derived PK model for both oral and implant dosing with the best-fit parameters in Table 1. All dose-dependent parameters are given in Suppl. Table S1. Corresponding model-simulated and super-imposed clinical pharmacokinetics are depicted in Fig. 1B–C.

Following rapid absorption, the plasma PK exhibits three phases: The initial phase involves drug distribution from plasma C to peripheral compartments. In this pre-steady state, the transfer rate to the second peripheral compartment C *→* P_2_ surpasses that of the first peripheral compartment C *→* P_1_. The second phase is dominated by C *→* P_1_. Subsequently, concentration exchange between compartments becomes constant, reaching a steady state. The slow elimination of plasma ISL in the steady state is dominated by the flux P_1_ *→* C, with further details in Supplementary Figure S2.

Intracellular pharmacokinetics are characterized by a rapid uptake and conversion of ISL to ISL-TP and a slow elimination of ISL-TP from the intracellular compartment.

For visual guidance, we depict the concentration that would prevent 90% of infections (EC_90_), as computed from Suppl. Figure S3. As shown in Fig. 1B–C, single oral doses of ISL smaller than 2 mg and implants with less than 56 mg ISL are not sufficient to produce intracellular concentrations that surpass EC_90_.

### Pharmacodynamics

As a next step, we used the PK model to estimate HIV-1 viral dynamics after single oral doses of ISL as mono-therapy. We coupled the model to an established viral dynamics model as outlined in section *Methods* and estimated the potency IC_50_ of ISL-TP in the intracellular compartment (see Supplementary Figure S4).

Parameter estimation yielded IC_50_ = 429.707 *±* 6.7 *·* 10*^−^*^5^ nM. Fig. 2 shows predicted viral load (VL) profiles, incorporating clinical VL dynamics, super-imposed for single oral doses of 0.5, 1, 2, 10, and 30 mg as 10-days monotherapy in HIV-infected individuals.

**Fig 2.**
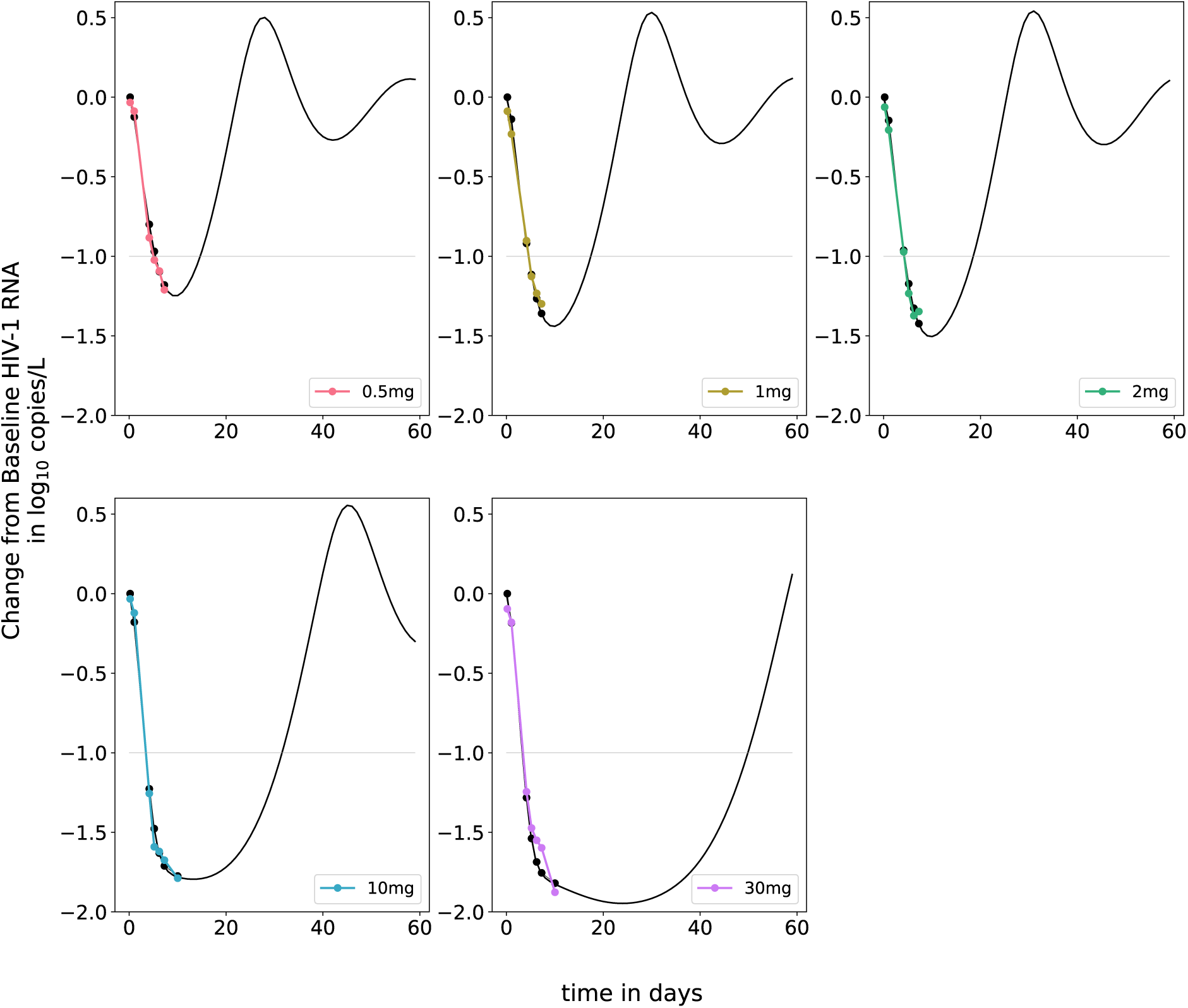
Pharmacodynamics of ISL. Clinical viral load reductions after a single oral dose of ISL (colored-dotted lines) vs. model-predicted viral load reductions (black-dotted lines). Data was derived from [19, 28]. The horizontal light gray line indicates a ten-fold virus load decline. Viral load data was used to estimate the potency of intracellular ISL-TP IC_50_, as described in the *Methods*.

All considered dosages result in a more than 10-fold VL decline by day 7 post-dose, as reported elsewhere [19, 28], whereby dose-depend declines can be observed both in the data and simulations. In the simulations, a single oral dose of 30 mg would result in a 100-fold decline in VL 23 days post-dosing.

### Predicted PrEP efficacy and toxicity for daily, weekly, biweekly and monthly oral dosing schemes

Recent work [39] using ISL phase II and III data highlighted that daily oral dosing of 0.25 mg ISL resulted in no observable adverse effects on lymphocytes and CD4^+^ T-cells counts. Assuming that plasma ISL is a correlate of toxicity, we found that a maximum plasma concentration of 7.02 nM is reached for daily oral dosing of 0.25 mg ISL. This value was used as a toxicity (NOAEL) threshold.

On the other hand, oral 0.75 mg once-daily led to observable decreases in lymphocytes, which were, however, reversible [38]. To evaluate putative toxicity of other daily, weekly and monthly dosing regimens, we computed the percentage of time (at steady state) where the plasma ISL concentration was above the NOAEL of 7.02 nM, as presented in Table 2. We only evaluated dosing regimens that generated plasma ISL levels either below the NOAEL or above the NOAEL for very few time points. Fig. 3 (top left) depicts simulated intracellular ISL-TP levels after 0.1 mg, 0.25 mg, and 0.5 mg once-daily dosing. As can be seen from the figure, a daily oral administration of ISL with 0.1 mg would provide a prophylactic efficacy of more than 90% 12 days after the first dose, while 0.25 mg (non-toxic) achieved *φ >* 90% after five dosing events and 0.5 mg after three dosing events. However, 0.5 mg may lead to lymphocyte drops, according to our ‘time-over-NOAEL’ analysis (Table 2).

**Fig 3.**
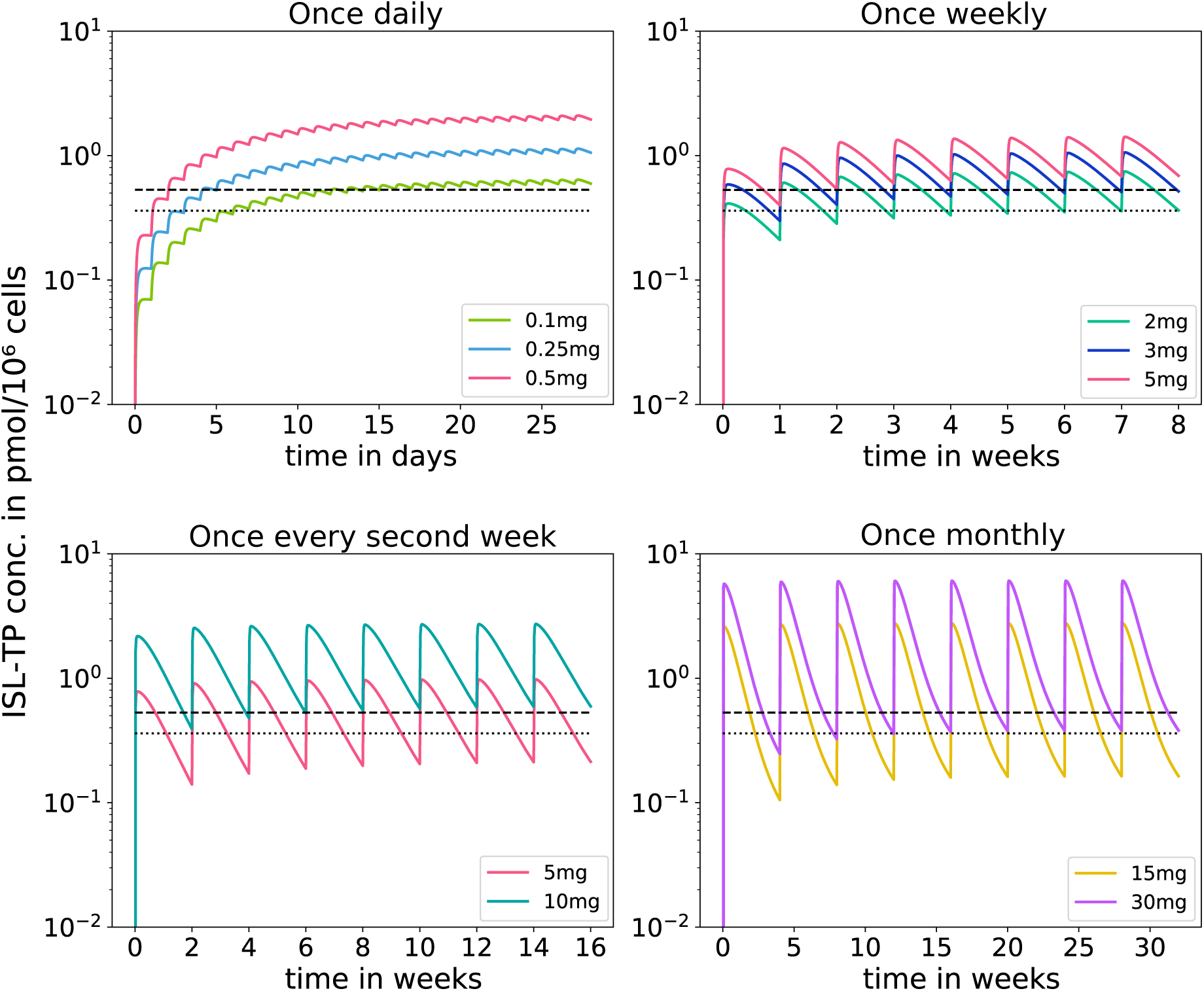
Predicted pharmacokinetics for once-daily, -weekly, -biweekly and -monthly oral ISL dosing schedules. Colored lines represent the simulated concentrations of different drug amounts for the respective dosing schedules, which are considered non-toxic (according to our findings in Table 2). The horizontal-dashed lines indicate model-computed 85- and 90% HIV-protective intracellular ISL-TP concentrations (EC_90_).

**Table 2.** Summary of derived efficacy vs. toxicity levels for all investigated regimen. The column ‘Time above threshold’ was used to determine the toxicological risks of the investigated regimen: It depicts the fraction of time that the plasma ISL concentration at steady state is above a no-observed-adverse-effect-level (NOAEL) of 7.02 nM, with a traffic-light related risk grading (green = safe, red = toxic).

| Dose in mg | Drug regimen | Dosing interval $\tau$ (in hrs) | Plasma AUC for $\tau = 24$ h | Time above threshold | $\phi \geq 90$ % | $\phi \geq 85$ % |
| --- | --- | --- | --- | --- | --- | --- |
| 0.1 | once daily | 24 | 12.99 | 0.0% | Yes | Yes |
| 0.25 | once daily | 24 | 32.49 | 0.0% | Yes | Yes |
| 0.5 | once daily | 24 | 64.97 | 9.5% | Yes | Yes |
| 0.75 | once daily | 24 | 97.46 | 14.9% | Yes | Yes |
| 2 | once weekly | 168 | 36.89 | 3.4% | No | Yes |
| 3 | once weekly | 168 | 55.33 | 4.2% | Yes | Yes |
| 5 | once weekly | 168 | 92.21 | 5.4% | Yes | Yes |
| 10 | once weekly | 168 | 184.44 | 9.7% | Yes | Yes |
| 15 | once weekly | 168 | 276.66 | 27.1% | Yes | Yes |
| 5 | once biweekly | 336 | 46.37 | 2.6% | No | No |
| 10 | once biweekly | 336 | 92.73 | 4.0% | Yes | Yes |
| 15 | once biweekly | 336 | 139.1 | 8.9% | Yes | Yes |
| 30 | once biweekly | 336 | 278.2 | 25.2% | Yes | Yes |
| 10 | once monthly | 672 | 46.22 | 1.9% | No | No |
| 15 | once monthly | 672 | 69.32 | 3.6% | No | No |
| 30 | once monthly | 672 | 138.65 | 10.8% | No | Yes |
| 60 | once monthly | 672 | 277.29 | 19.4% | No | Yes |
| 1 | on demand (1-0-0) | 120 | 20.25 | 2.8% | No | No |
| 2 | on demand (1-0-0) | 120 | 40.5 | 4.4% | No | Yes |
| 5 | on demand (1-0-0) | 120 | 101.24 | 7.6% | Yes | Yes |
| 10 | on demand (1-0-0) | 120 | 202.47 | 10.3% | Yes | Yes |
| 0.25 | on demand (2-1-1) | 192 | 13.19 | 0.9% | No | Yes |
| 0.5 | on demand (2-1-1) | 192 | 26.37 | 3.8% | Yes | Yes |
| 0.75 | on demand (2-1-1) | 192 | 39.56 | 5.5% | Yes | Yes |
| 1 | on demand (2-1-1) | 192 | 52.75 | 6.9% | Yes | Yes |
| 2 | on demand (2-1-1) | 192 | 105.49 | 10.4% | Yes | Yes |
| 5 | on demand (2-1-1) | 192 | 263.72 | 20.7% | Yes | Yes |
| 48 | Implant | 2016 | 6.51 | 0.0% | No | No |
| 52 | Implant | 2016 | 9.2 | 0.0% | No | No |
| 54 | Implant | 2016 | 10.27 | 0.1% | No | No |
| 56 | Implant | 2016 | 11.83 | 0.2% | Yes | Yes |
| 62 | Implant | 2016 | 15.71 | 0.2% | Yes | Yes |

If taken once-weekly at least 5 mg are necessary to achieve *>* 90% protection. However, this dosing regimen is associated with 5% of time over NOAEL. If ISL is taken every second week, 10 mg would be required, with 4% of time over NOAEL. For monthly dosing, at least 30 mg need to be taken, but ISL-TP trough levels may fall under the 90% protective threshold. Moreover, the monthly dosing regimen would be associated with substantial time over NOAEL (Table 2).

### Prophylactic efficacy and toxicity of an on-demand regimen around the time of exposure

Established on-demand PrEP [9, 40] dosing patterns with FTC/TDF for MSM consist of a double dose followed by two pills on consecutive days (2 *−* 1 *−* 1). In our simulations, we also investigated the efficacy profile of low single oral doses (1 *−* ∅ *−* ∅).

Fig. 4A, shows prophylactic efficacy profiles when viral challenge occurred shortly before (PEP), or after (PrEP) the first dose at time point *t* = 0 in the on-demand regimens. The time points at which a regimen falls below an efficacy threshold are shown by black dashed vertical lines. Notably, an on-demand regimen could circumvent toxic effects, since plasma ISL may only stay above NOAEL for very short times, or does not even reach NOAEL.

**Fig 4.**
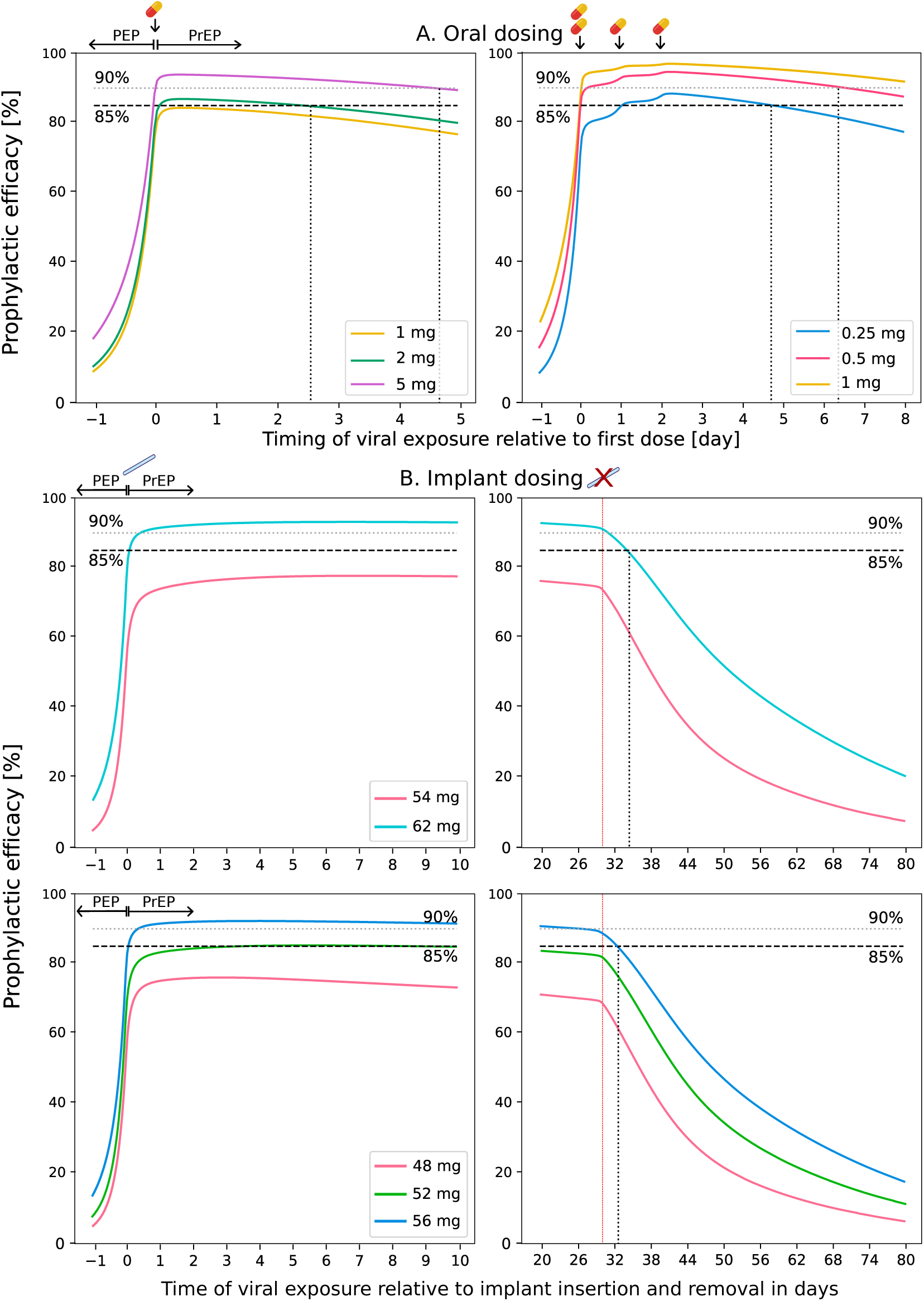
Prophylactic efficacy of oral on-demand dosing and implant administration around the time of virus exposure. A. Prophylactic efficacy of oral 1 *−* ∅ *−* ∅ and 2 *−* 1 *−* 1 dosing schemes around the time of virus exposure (indicated on the x-axis). B. Prophylactic efficacy of subdermal implants shortly after insertion and -removal. The removal of the implants is shown as red vertical lines. The x-axis indicates the time of virus exposure, relative to the first dosing event (or implant insertion event). Prophylactic efficacy values of *φ* = 85% and *φ* = 90% are depicted as dashed and dotted horizonal lines, respectively. Vertical black dotted lines indicate the time, when a considered regimen falls below an efficacy threshold.

According to our simulations, a single ingestion of 2 mg oral ISL would give rise to *>* 85% prophylactic efficacy almost immediately if taken at the same time as virus exposure or up to 2.5 days prior to the exposure event but never reaches 90%. A single oral 5 mg tablet ISL would reach protective levels of *>* 90% similarly fast after dosing but would remain protective for about 5 days. Both dosing regimens would give rise to plasma ISL levels that are above NOAEL for 4.4% and 7.6% of time, see Table 2.

For the 2 *−* 1 *−* 1 regimen, only oral doses of 0.5, 1 mg or above would yield *>* 90% efficacy. For 0.5 mg, this level of protection is only achieved if virus exposure occurred at least 3 hours after the first dose. A prophylactic efficacy *>* 90% is maintained for 6 vs. more than 8 days after the last dose with 0.5 and 1 mg, respectively, during 2 *−* 1 *−* 1 on-demand dosing. However, both dosages lead to substantial time (3.8% vs. 6.9%) above NOAEL (Table 2). Furthermore, we observe that all considered dosages are too low to provide sufficient post-exposure prophylactic (PEP) efficacy, while larger dosages could cause adverse effects and were therefore not further considered.

### Prophylactic efficacy and toxicity of ISL implants

We simulated the prophylactic efficacy of implants loaded with 48, 52, 54, 56 and 62 mg with emphasis on the time shortly after implant insertion (’how quickly is protection achieved?’) and after implant removal, Fig. 4B. According to our simulations, all considered drug loads lead to plasma ISL exposures smaller than the NOAEL, Table 2. Drug loads greater than 56 mg ISL would give rise to *>* 90% protection as early as a few hours after implant insertion. After implant removal, the loading dose of 62 mg falls below the 90% efficacy threshold after about 1.5 days and below the 85% threshold after more than 4 days.

## Discussion

Adherence to daily oral PrEP is challenging, limiting its uptake. Injectable CAB-LA may overcome daily dosing requirements, but the necessity of frequent injections along with sub-optimal drug access may constrain its global impact on HIV prevention. The availability of additional options could contribute to satisfying unmeet PrEP-needs. In this work, we identified ISL implant regimens that may be a safe and effective against HIV transmission.

We examined the PK/PD properties of ISL using an integrated mathematical model that represents two dosing routes (oral and subdermal implant). It comprises three compartments with an additional intracellular effect compartment (Fig. 1A). According to our model, the ISL concentration-time profile in plasma exhibits three phases: In the first phase, the drug concentration enters the plasma and partially diffuses into the peripheral compartments (C *→* P_2_ *>* C *→* P_1_). The second phase is characterized by the flux into the first peripheral compartment (C *→* P_1_). The final phase is a quasi-steady state where distribution reaches equilibrium, and plasma ISL elimination is primarily dominated by the rate-limiting flux from P_1_ *→* C (Supplementary Figure S2).

We observed dose-proportionality in plasma PK for all considered doses (0.5–400 mg, Fig. 1C), consistent with prior findings [18]. Implant dosing resulted in an initial sharp decline of plasma concentrations, transitioning into a very slow decay after about 24 h. The sharp decline was best modeled by an initial drug concentration in the subcutaneous compartment after implant insertion (details see *Methods, PK modeling*), possibly arising from minor injuries at the implant insertion site. While plasma ISL concentrations for the subdermal implant were associated with dose based on available data, they may not necessarily be dose-proportional.

Intracellular ISL-TP did not show proportionality to plasma ISL at higher oral doses (*≥* 30 mg, Supplementary Table S1). However, we constructed a non-linear function that modeled the relationship between cellular uptake, intracellular anabolism (*k_C→E_*), and plasma concentrations for oral dosing (Supplementary Figure S5) to interpolate between oral dosing regimens. The active moiety ISL-TP had an intracellular half-life of 87.75 hours in our model, aligning with literature findings [19] of 78.5–128 h for oral dosages of 0.5 mg to 30 mg. Its long half-life and high potency allows for sparse dosing frequencies as evaluated in Fig. 3. To validate our PK model, we simulated two oral regimens that were not included in parameter estimation, Supplementary Figure S1 showing overall satisfactory match, particularly for the low-dose regimen, which were the focus of this study.

For implants, intracellular ISL-TP concentration data was not directly correlated with drug loading. The available data showed that the ISL-TP concentration for 52 mg exceeded that of 54 mg (Fig. 1B). However, this observation may be linked to an experimental artifact where ISL-TP concentrations for implants might be under-reported (potentially higher than experimentally determined). In our evaluations of prophylactic efficacy, 52 mg and 54 mg resulted in suboptimal HIV protection (*<* 90%) and were not considered further. This observation might be a conservative estimate for 54 mg if the data under-reports ISL-TP levels with this regimen.

Following PK model construction and parameterization, we used clinical phase-II data to estimate the remaining free parameter (antiviral potency IC_50_ of ISL-TP). Specifically, we used VL data following single oral doses of 0.5 mg to 30 mg ISL reported in [28] and estimated IC_50_ = 429.707 nM by fitting model predicted and reported VL data, Fig. 2.

Recently, it was reported that once-daily administration of 0.75 mg ISL leads to a reversible reduction in lymphocyte counts compared to other antiretroviral drugs [41]. While historically, the inhibition of human DNA polymerases (in particular mitochondrial polymerase *γ*) is associated with NRTI toxicity [15, 42], ISL does not inhibit human DNA polymerase *γ* [41]. Sequential intracellular phosphorylation by cellular kinases is essential for the biological activation of the drug within HIV target cells. The same enzymes are also used to phosphorylate endogenous nucleosides required for cell metabolism and proliferation. In line with recent discussions on potential mechanisms of NRTI toxicity, the toxic effect of ISL may be due to competition (and inhibition) with endogenous nucleosides for phosphorylation in lymphocytes [43–45]. Notably, a similar mechanism was observed regarding the cardiac toxicity of the NRTI zidovudine (AZT) [46] and denotes the mechanistic basis of synergy between the NRTIs tenofovir and emtricitabine [16]. In both examples, the effect strength (toxicity, synergy) correlates with plasma pro-drug concentrations. Since ISL is a high-affinity substrate for import and phosphorylation, it may be reasonable to assume that a metabolic predecessor of ISL-TP interferes with cellular (deoxy-)nucleo(s/t)ide pools as causative mechanism of lymphopenia [47]. Importantly, in this case, not ISL-TP, but rather metabolic predecessors may correlate with toxicity. Thus, we assumed that plasma ISL is a correlate of toxicity since the exact mechanisms behind the reduction of lymphocytes are still unknown to date.

To evaluate the putative toxicity of ISL, we replicated dosing scenarios previously examined for toxic endpoints (lymphocyte count reduction) [38, 39] to identify maximal ISL concentrations that do not cause adverse effects. At 0.25 mg once-daily, we identified a NOAEL threshold of 7.02 nM. Subsequently, we determined the fraction of time, for a given dosing schedule, during which plasma ISL exceeded the NOAEL. This approach indicates a very strict (cautious, conservative) assessment of potential toxicity. An alternative assessment based on the AUC over a 24 h dosing interval is presented in Table 2.

Based on the integrated PK-PD-viral dynamics model, we performed simulations to evaluate prophylactic efficacy after oral or implant dosing. For these simulations, we focused on regimens that would achieve sufficient protection (*φ >* 85% and *φ >* 90%) and minimize risks of lymphopenia, as outlined above.

Simulation of once-daily oral ISL at non-toxic doses of 0.1 and 0.25 mg achieves 90% prophylactic efficacy, with the 0.25 mg regimen requiring less than daily adherence (4/7 pills achieve *φ >* 90%, see Supplementary Figure S6). Compared to established oral PrEP with FTC/TDF, the adherence-efficacy profile of 0.25 mg may be slightly inferior [10, 48, 49] and does not offer a unique niche in the prophylactic portfolio. Less frequent dosing schedules (weekly, bi-weekly, monthly) achieve *>* 90% protection but entail a residual risk of toxic side effects. On-demand regimens for PrEP/PEP scenarios suggest no effective protection without potential side effects, according to our conservative estimates.

In this evaluation, ISL-TP in PBMCs served as an efficacy marker for HIV PrEP, demonstrating proportional kinetics to exposed tissues (vaginal and rectal) after sexual HIV challenges [50, 51]. While PBMCs contain a significant portion of HIV target cells (CD4^+^ T-cells) that are decisive for establishing viral infection, tissue samples primarily consist of non-HIV target cells (e.g. epithelial cells) with minimal tissue-resident CD4^+^ T-cells. Since NRT(T)Is undergo active transport and intracellular conversion by kinases, which may vary in cell-type-specific expression levels, analyzing tissue samples and cell mixtures dominated by cells irrelevant to viral replication may not accurately predict efficacy for this drug class. Consistent with these considerations, recent findings suggest that PBMCs serve as a better efficacy marker for oral FTC/TDF-based PrEP compared to local tissue concentrations in case of sexual HIV challenge [10].

Our modeling indicates that subdermal ISL implants with at least 56 mg are effective and safe for LA-PrEP, providing prophylactic efficacy while minimizing toxicity risks (0.2% according to our criteria, Table 2). However, the putative toxicity mechanisms and markers for ISL and NRTTIs require further elucidation. The proposed model could be adapted for novel NRTTIs, including Merck’s new developmental drug NCT05494736 (MK-8527), to support clinical investigation and identify prophylactic niches [52].

## Study highlights

### What is the current knowledge on the topic?

Islatravir (ISL) is an extremely potent first-in-class nucleoside-reverse-transcriptase-translocation-inhibitor (NRTTI) that may be suitable for infrequent (i.e. weekly) oral dosing as HIV pre-exposure prophylaxis (PrEP), or as long-acting (LA-)PrEP when administered as subdermal implant. However, lymphopenia was associated with oral 0.75 mg daily, putting PrEP trials on halt and potentially limiting treatment use, although further research is ongoing in this area.

### What question did this study address?

We modeled the pharmacokinetics, pharmacodynamics, toxicity and prophylactic efficacy of ISL to assess the safety and suitability of low oral doses or implant dosing for HIV prevention.

### What this study adds to our knowledge?

Our findings suggest that implants with 56–62 mg ISL offer safe, effective HIV risk reduction without risk of lymphopenia. Low doses of oral ISL can be safe but constitute adherence requirements similar to established oral PrEP regimen.

### How might this change drug discovery, development, and/or therapeutics?

Our work highlights that oral ISL at effective, non-toxic doses may not fill a unique prophylactic niche, while implant dosing could be further explored as LA-PrEP. Our modeling may serve as a blueprint for defining the prophylactic niche of second-generation NRTTIs.

## Acknowledgement

H.K., L.Z. and M.v.K. acknowledge funding from the German Ministry for Science and Education (BMBF; grants 01KI2016). The funders had no role in designing the research or the decision to publish.

## Conflict of interest statement

J.H. has been a consultant for Merck; prior studies have received donated FTC-TDF from Gilead. C.W.H. has received funding for clinical research and been a (past) consultant for Merck, Gilead and ViiV/GSK, holds US patents related to HIV prevention technology (US20200138700A1) and founded Prionde Biopharma, LLC, a microbicide company; all relationships are managed by Johns Hopkins University.

## Author Contributions

H.K. and M.v.K. wrote the manuscript with help from C.W.H. and J.H. H.K. and M.v.K. designed the research. H.K. and L.Z. performed the research and H.K., C.W.H., J.H. and M.v.K analyzed the data.

## 1 Supporting Information

**Table S1.** Dosage-specific PK parameters. Rate constants are given in the units *h^−^*^1^ except the implant-specific initial concentrations which is denotes the percentage of the loading dose ld. Drug release rates *k*_Impl_*_→_*_SC_(ld) for loading doses ld = 54 and 62 mg implants were determined using cumulative drug release profiles for non-erodible EVA from [31]. Release rates for loading doses of 48, 52, and 56 mg ISL were obtained by linear interpolation. Parameter *k_R_* (associated with the drug release) is global for all dosages. The initial drug concentrations for the subcutaneous compartment *π*_ld_ was set to *π*_54_ if the drug load was smaller or equal to 54 mg and to *π*_62_ otherwise. Parameter *π*_ld_ could only be determined for loading doses of 54 mg and 62 mg, since plasma ISL was only available for these loading doses (see *Methods*). Intracellular *k_C→E_* values are given in mean *±* bootstrap-estimated standard deviation.

| Parameter | Value | Parameter | Value |
| --- | --- | --- | --- |
| $k_R$ | $2.18 \cdot 10^{-4}$ (global) | $\pi_{48}$ | 4.12 % (48 mg) |
| $k_{\text{Impl} \rightarrow \text{SC}}(48)$ | $1.92 \cdot 10^{-4}$ | $\pi_{52}$ | 4.12 % (52 mg) |
| $k_{\text{Impl} \rightarrow \text{SC}}(52)$ | $2.77 \cdot 10^{-4}$ | $\pi_{54}$ | 4.12 % (54 mg) |
| $k_{\text{Impl} \rightarrow \text{SC}}(54)$ | $3.07 \cdot 10^{-4}$ | $\pi_{56}$ | 5.83 % (56 mg) |
| $k_{\text{Impl} \rightarrow \text{SC}}(56)$ | $3.45 \cdot 10^{-4}$ | $\pi_{62}$ | 5.83 % (62 mg) |
| $k_{\text{Impl} \rightarrow \text{SC}}(62)$ | $4.60 \cdot 10^{-4}$ | | |

| Dose in mg | $k_{C \rightarrow E}$ (oral) | Dose in mg | $k_{C \rightarrow E}$ (implant) |
| --- | --- | --- | --- |
| 0.5 | $37.69 \pm 1.4437$ | 48 | $25.30 \pm 0.7982$ |
| 1 | $28.17 \pm 1.0711$ | 52 | $32.55 \pm 1.0794$ |
| 2 | $16.56 \pm 0.5626$ | 54 | $19.35 \pm 0.7048$ |
| 5 | $12.59 \pm 0.6236$ | 56 | $41.98 \pm 1.4276$ |
| 10 | $17.46 \pm 0.5703$ | 62 | $36.87 \pm 1.1198$ |
| 15 | $13.81 \pm 1.0272$ | | |
| 30 | $16.95 \pm 0.4831$ | | |
| 100 | $08.83 \pm 0.4775$ | | |
| 200 | $10.57 \pm 0.5794$ | | |
| 400 | $09.10 \pm 0.5627$ | | |

**Figure S1.**
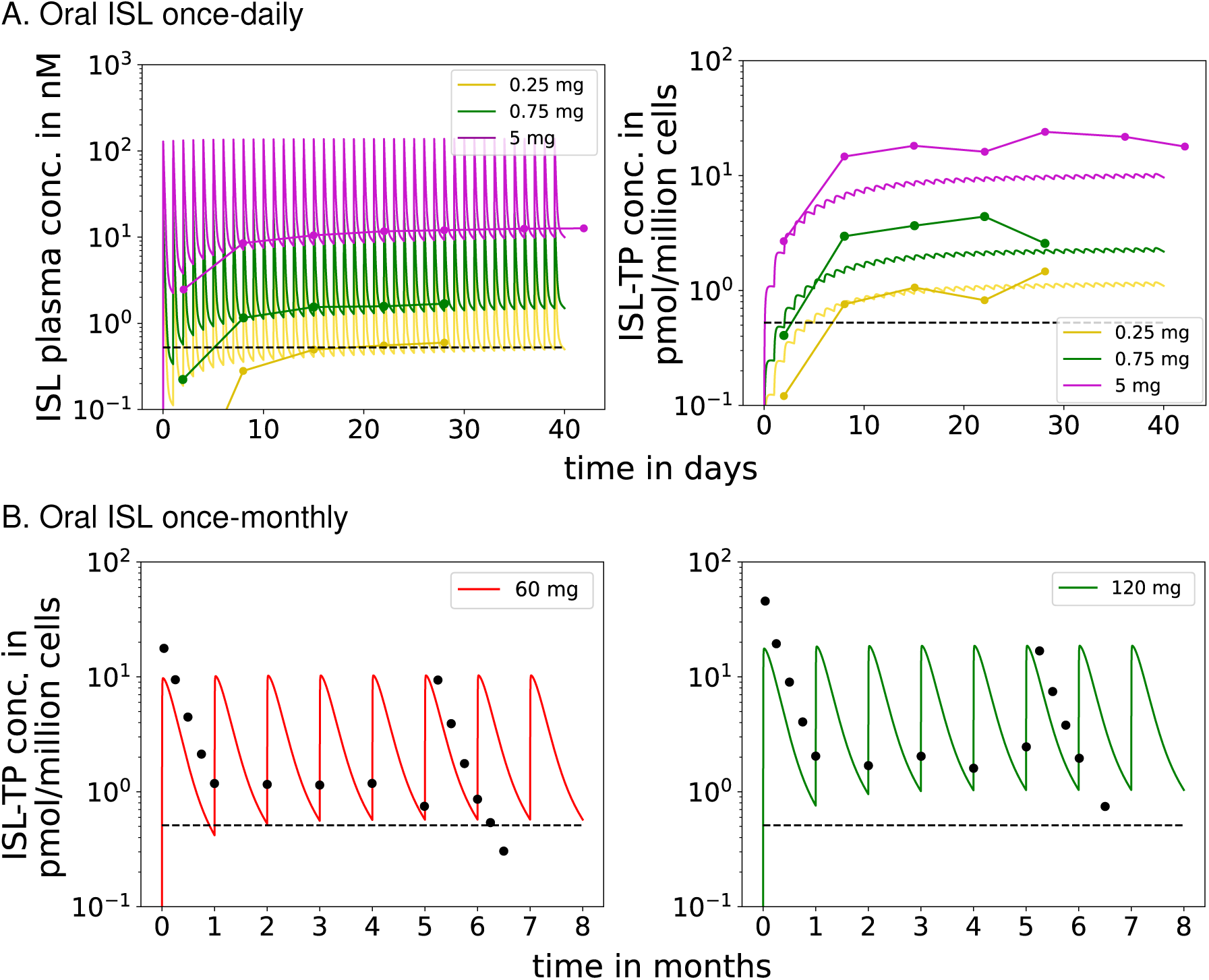
Cross-Validation of the PK model with independent data set. A. Model-predicted (continous lines) and data derived [36] mean ISL plasma and intracellular ISL-TP trough concentrations (lines with dots, *n* = 9 participants per dose) after daily oral doses of 0.25, 0.75 and 5 mg ISL in individuals without HIV infection. B. Model-predicted (solid lines) and data derived [37] mean intracellular ISL-TP concentrations (black dots) after once-monthly oral doses of 60 mg and 120 mg ISL. The datasets were not used in parameter estimation and just served as a cross-validation of the model.

**Figure S2.**
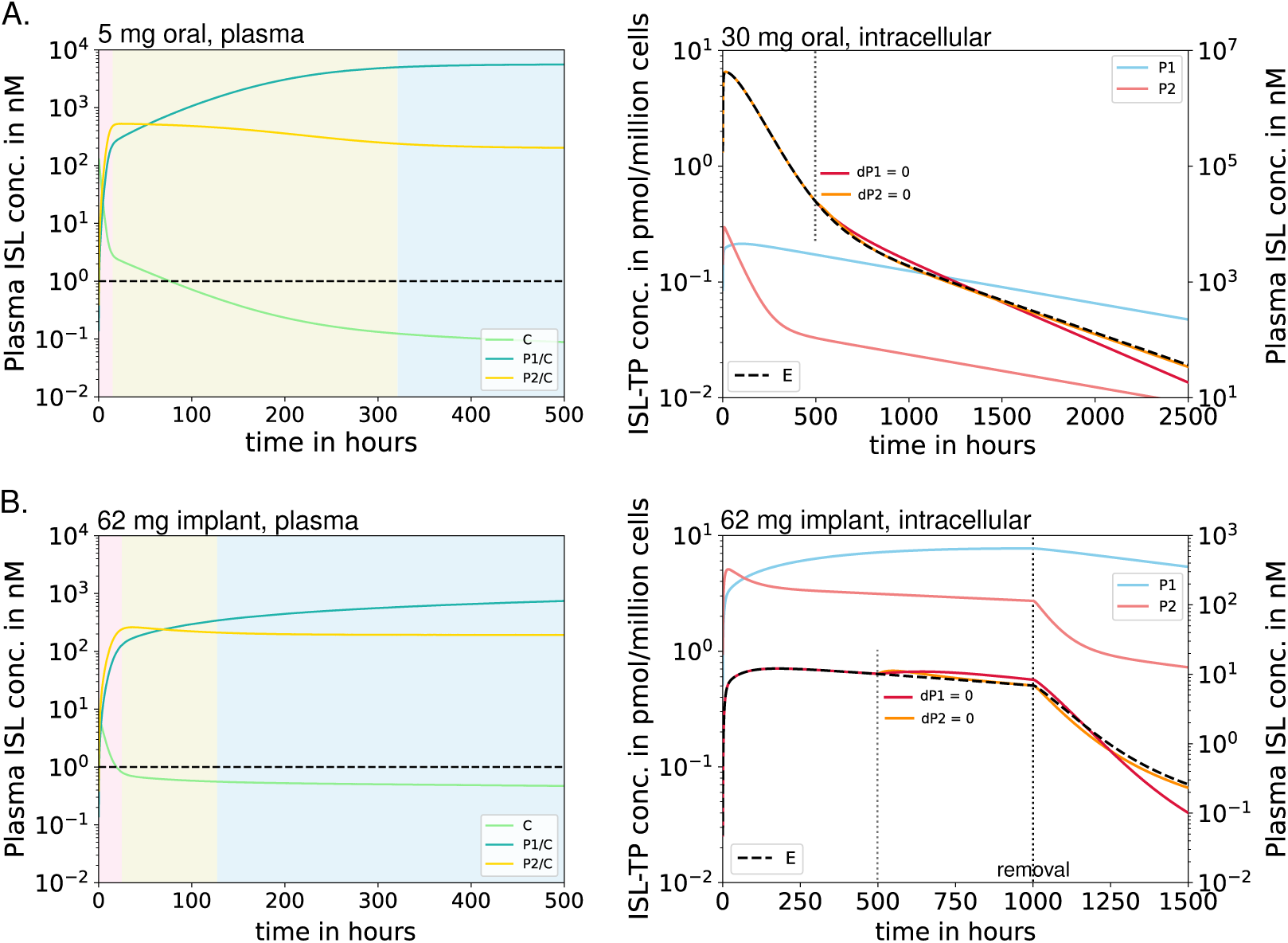
Three-phasic ISL decay. The figure illustrates concentration-time profiles of ISL in plasma *C* and intracellular ISL-TP (E) following a single oral dose (panel A) vs. implant dosing and removal at *t* = 1000 hours (panel B). On the left, the three concentration distribution phases, illustrated as colored areas, are shown, encompassing two pre-steady-state phases followed by a final steady-state phase: (i) The drug enters the plasma and is distributed in peripheral compartments, with C *→* P_2_ *>* C *→* P_1_ (indicated by an uptake into P_2_ (yellow line) which is reflected by a decay in plasma ISL (green line). (ii) Since the second peripheral compartment P_2_ is saturated faster than the first, the second phase shows mainly the concentration flow from the plasma into the first peripheral compartment (C *→* P_1_). I.e. the decay in plasma ISL (green line) is reflected by an increase in P_1_ (blue line). (iii) The final phase reflects equilibrium in all compartments where P_1_ *→* C dominates (details on the right). Right-side figures show the impact of peripheral compartments (P_1_=light blue, P_2_=light pink; right y-axis) on the intracellular ISL-TP pharmacokinetics (compartment E, dashed line, left y-axis). As a test, we either “switched off” the first peripheral compartment (*d*P_1_ = 0) at *t* = 500 h, or the second peripheral compartment (*d*P_2_ = 0) and observed the pharmacokinetics of intracellular ISL-TP, red vs. orange line respectively. Switching off a compartment involves preventing the concentration from entering the corresponding compartment while still allowing transfer to the central compartment. When the second peripheral compartment is “switched off”, we observe no difference in intracellular ISL-TP in comparison to the full model (dashed black line vs. orange line). In contrast, if the first peripheral compartment is “switched off”, there is a more rapid decay of ISL-TP in the final phase. This indicates that P_1_ acts as a reservoir for the final decay phase of intracellular ISL-TP, while P_2_ has no impact on the final decay phase (but does impact the initial decline).

**Figure S3.**
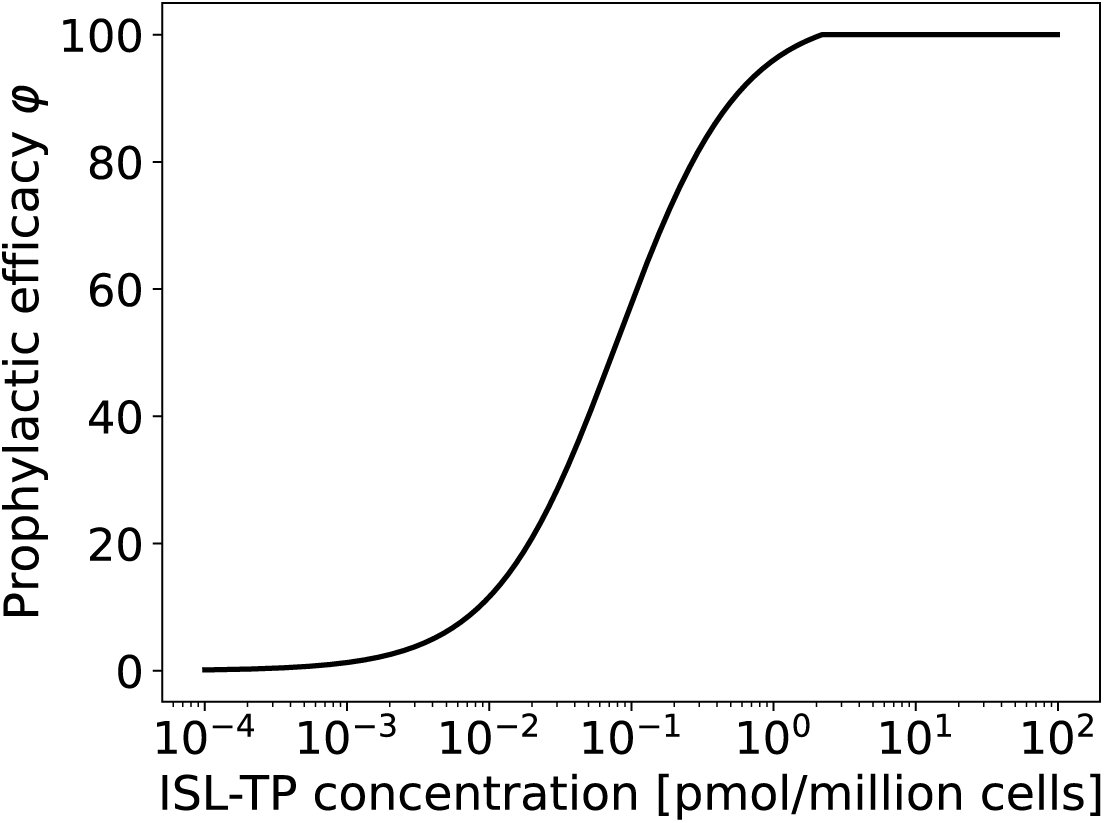
Derivation of prophylactic efficacy thresholds. Prophylactic efficacy for a range of constant intracellular ISL-TP concentrations was computed analytically using the methods in [53]. With a direct effect IC_50_ of 429.707 nM (inhibition of reverse transcription; fitted to viral dynamics), we obtained 85% and 90% prophylactic efficacy (HIV risk reduction) thresholds EC_85_ = 0.35 pmol/million cells and EC_90_ = 0.51 pmol/million cells in PBMCs, respectively.

**Figure S4.**
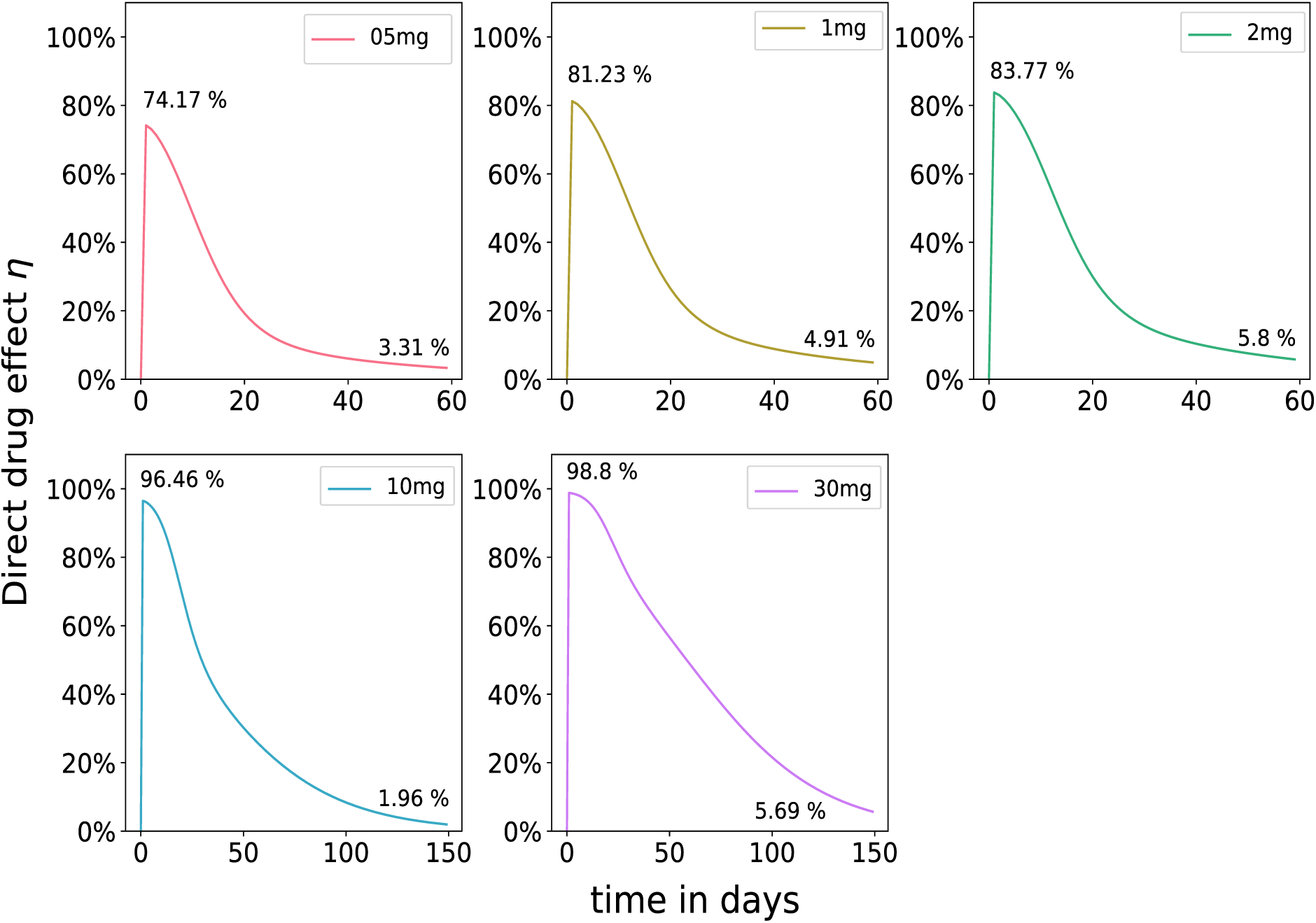
Simualated direct drug effects of ISL after a single oral dose. Direct effect *η* (reduction in reverse transcription; Eq. 8, see *Methods*) after a single oral dose of ISL. Colored lines represent the respective dosages ranging from 0.5 mg to 30 mg.

**Figure S5.**
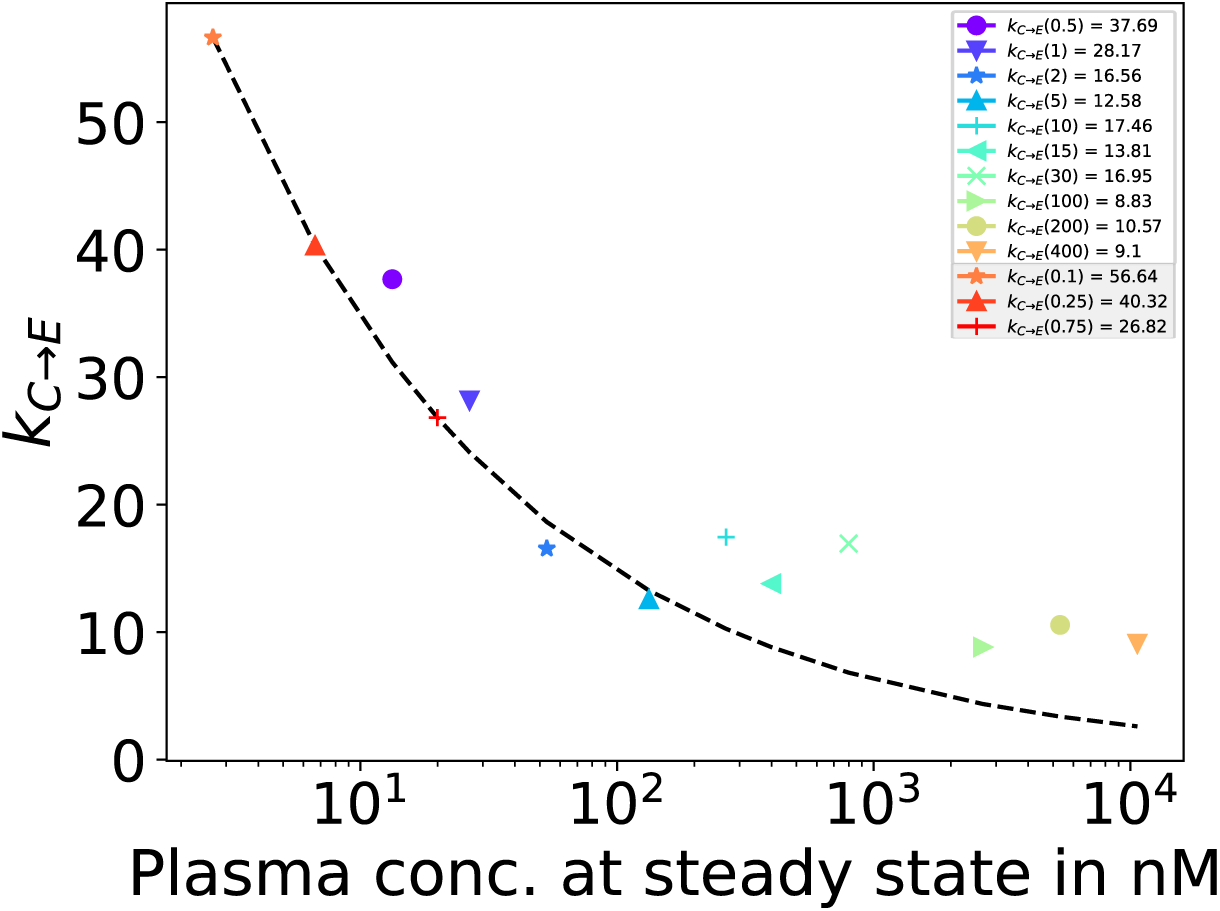
Extrapolation of dose-dependent parameter. *k_C→E_*. Relation of *k_C→E_*(rate constant for ISL uptake and intracellular conversion) to plasma concentrations *x* at steady state for oral data with weekly (168 h) dosing events. Parameter *k_C→E_* was fitted from plasma data for all dosages indicated in Fig 1. These fitted parameters could be approximated by the function *k_C→E_*= *x^−^*^0.371^ *· e*^4.4^. Since oral doses of 0.1, 0,25 and 0.75 mg were not contained in the data used for parameter estimation, dose-dependent parameters *k_C→E_*for these dosages were determined using the function above (red star, triangle and cross).

**Figure S6.**
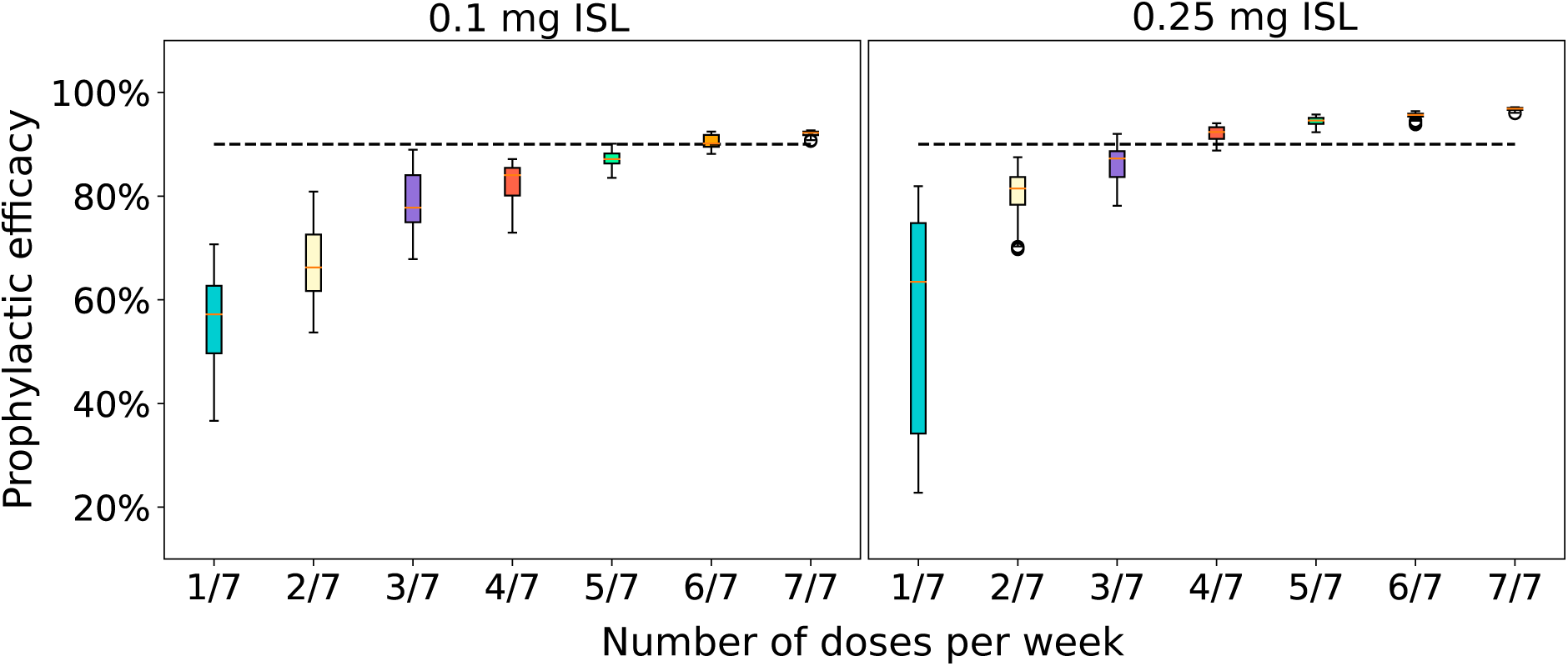
Adherence requirements for oral PrEP. We simulated oral doses of 0.1 mg (left) and 0.25 mg (right) ISL with varying levels of adherence (x-axis) and computed the resulting prophylactic efficacy. Boxplots show the median efficacy, the interquartile range and whiskers extend to 3 *×* the interquartile range.

## References

1. F. Barré-Sinoussi, J. C. Chermann, F. Rey, M. T. Nugeyre, S. Chamaret, J. Gruest, C. Dauguet, C. Axler-Blin, F. Vézinet-Brun, C. Rouzioux, W. Rozenbaum, and L. Montagnier. Isolation of a T-Lymphotropic Retrovirus from a Patient at Risk for Acquired Immune Deficiency Syndrome (AIDS). Science, 220(4599):868–871, may 1983.

2 Robert C Gallo. Virus hunting: AIDS, cancer, and the human retrovirus: A story of scientific discovery. BasicBooks, 1991.

3. Fact sheet - Latest global and regional statistics on the status of the AIDS epidemic. https://www.unaids.org/sites/default/files/media_asset/UNAIDS_FactSheet_en.pdf. Accessed on Fri, January 06, 2023.

4. UNAIDS Global AIDS Update 2022 https://www.unaids.org/sites/default/files/media_asset/2022-global-aids-update-summary_en.pdf. Accessed on Wed, January 04, 2023.

5. Robert M Grant, Javier R Lama, Peter L Anderson, Vanessa McMahan, Albert Y Liu, Lorena Vargas, Pedro Goicochea, Martín Casapía, Juan Vicente Guanira-Carranza, Maria E Ramirez-Cardich, et al. Preexposure chemoprophylaxis for HIV prevention in men who have sex with men. New England Journal of Medicine, 363(27):2587–2599, 2010.

6. Jared M Baeten, Thesla Palanee-Phillips, Elizabeth R Brown, Katie Schwartz, Lydia E Soto-Torres, Vaneshree Govender, Nyaradzo M Mgodi, Flavia Matovu Kiweewa, Gonasagrie Nair, Felix Mhlanga, et al. Use of a vaginal ring containing dapivirine for hiv-1 prevention in women. New England Journal of Medicine, 375(22):2121–2132, 2016.

7. Daniel Schmidt, Yannick Duport, Christian Kollan, Ulrich Marcus, Sara Iannuzzi, and Max von Kleist. Dynamics of HIV Prep Use and Coverage During and after Covid-19 in Germany. Available at SSRN: 10.2139/ssrn.4611490, 2023.

8. Jean-Michel Molina, Catherine Capitant, Bruno Spire, Gilles Pialoux, Laurent Cotte, Isabelle Charreau, Cecile Tremblay, Jean-Marie Le Gall, Eric Cua, Armelle Pasquet, et al. On-demand preexposure prophylaxis in men at high risk for HIV-1 infection. New England Journal of Medicine, 373(23):2237–2246, 2015.

9. Jean-Michel Molina, Jade Ghosn, Lambert Assoumou, Constance Delaugerre, Michéle Algarte-Genin, Gilles Pialoux, Christine Katlama, Laurence Slama, Geoffroy Liegeon, Lydie Beniguel, et al. Daily and on-demand HIV pre-exposure prophylaxis with emtricitabine and tenofovir disoproxil (ANRS PREVENIR): a prospective observational cohort study. The Lancet HIV, 9(8):e554–e562, 2022.

10. Lanxin Zhang, Sara Iannuzzi, Ayyappa Chaturvedula, Elizabeth Irungu, Jessica E Haberer, Craig W Hendrix, and Max von Kleist. Model-based predictions of protective hiv pre-exposure prophylaxis adherence levels in cisgender women. Nature medicine, pages 1–10, 2023.

11. World Health Organization and others. Differentiated and simplified pre-exposure prophylaxis for HIV prevention: update to WHO implementation guidance: technical brief. WHO, 2022.

12. Sinead Delany-Moretlwe, James P Hughes, Peter Bock, Samuel Gurrion Ouma, Portia Hunidzarira, Dishiki Kalonji, Noel Kayange, Joseph Makhema, Patricia Mandima, Carrie Mathew, et al. Cabotegravir for the prevention of HIV-1 in women: results from HPTN 084, a phase 3, randomised clinical trial. The Lancet, 399(10337):1779–1789, 2022.

13. Tracy L Diamond, Winnie Ngo, Min Xu, Shih Lin Goh, Silveria Rodriguez, Ming-Tain Lai, Ernest Asante-Appiah, and Jay A Grobler. Islatravir has a high barrier to resistance and exhibits a differentiated resistance profile from approved nucleoside reverse transcriptase inhibitors (NRTIs). Antimicrobial Agents and Chemotherapy, 66(6):e00133–22, 2022.

14. Fernanda P Pons-Faudoa, Nicola Di Trani, Simone Capuani, Jocelyn Nikita Campa-Carranza, Bharti Nehete, Suman Sharma, Kathryn A Shelton, Lane R Bushman, Farah Abdelmawla, Martin Williams, et al. Long-acting refillable nanofluidic implant confers protection against shiv infection in nonhuman primates. Science translational medicine, 15(702):eadg2887, 2023.

15. Max von Kleist, Philipp Metzner, Roland Marquet, and Christof Schütte. HIV-1 polymerase inhibition by nucleoside analogs: cellular-and kinetic parameters of efficacy, susceptibility and resistance selection. PLoS Computational Biology, 8(1):e1002359, 2012.

16. Sara Iannuzzi and Max von Kleist. Mathematical modelling of the molecular mechanisms of interaction of tenofovir with emtricitabine against HIV. Viruses, 13(7):1354, 2021.

17. Eleftherios Michailidis, Andrew D. Huber, Emily M. Ryan, Yee T. Ong, Maxwell D. Leslie, Kayla B. Matzek, Kamalendra Singh, Bruno Marchand, Ariel N. Hagedorn, Karen A. Kirby, Lisa C. Rohan, Eiichi N. Kodama, Hiroaki Mitsuya, Michael A. Parniak, and Stefan G. Sarafianos. 4*′*-Ethynyl-2-fluoro-2*′*-deoxyadenosine (EFdA) Inhibits HIV-1 Reverse Transcriptase with Multiple Mechanisms. Journal of Biological Chemistry, 289(35):24533–24548, aug 2014.

18. Randolph P Matthews, Wendy Ankrom, Evan Friedman, Deanne Jackson Rudd, Yang Liu, Robin Mogg, Deborah Panebianco, Inge De Lepeleire, Magdalena Petkova, Jay A Grobler, et al. Safety, tolerability, and pharmacokinetics of single-and multiple-dose administration of islatravir (mk-8591) in adults without hiv. Clinical and Translational Science, 14(5):1935–1944, 2021.

19. Dirk Schürmann, Deanne Jackson Rudd, Saijuan Zhang, Inge De Lepeleire, Martine Robberechts, Evan Friedman, Christian Keicher, Andreas Hüser, Jörg Hofmann, Jay A Grobler, S Aubrey Stoch, Marian Iwamoto, and Randolph P Matthews. Safety pharmacokinetics, and antiretroviral activity of islatravir (ISL MK-8591), a novel nucleoside reverse transcriptase translocation inhibitor, following single-dose administration to treatment-naive adults infected with HIV-1: an open-label, phase 1b, consecutive-panel trial. The Lancet HIV, 7(3):e164–e172, mar 2020.

20. Matthews, Randolph P, Zang, Xiaowei, Barrett, Stephanie E, Koynov, Athanas, Goodey, Adrian, Heimbach, Tycho, Weissler, Vanessa L, Leyssens, Carlien, Reynders, Tom, Xu, Zhiqing, et al. A randomized, double-blind, placebo-controlled, phase 1 trial of radiopaque islatravir-eluting subdermal implants for pre-exposure prophylaxis against hiv-1 infection. Journal of Acquired Immune Deficiency Syndromes (1999), 92(4):310, 2023.

21. Karen A Kirby, Eleftherios Michailidis, Tracy L Fetterly, Musetta A Steinbach, Kamalendra Singh, Bruno Marchand, Maxwell D Leslie, Ariel N Hagedorn, Eiichi N Kodama, Victor E Marquez, et al. Effects of substitutions at the 4’ and 2 positions on the bioactivity of 4’-ethynyl-2-fluoro-2’-deoxyadenosine. Antimicrobial agents and chemotherapy, 57(12):6254–6264, 2013.

22. Zhe Li Salie, Karen A Kirby, Eleftherios Michailidis, Bruno Marchand, Kamalendra Singh, Lisa C Rohan, Eiichi N Kodama, Hiroaki Mitsuya, Michael A Parniak, and Stefan G Sarafianos. Structural basis of HIV inhibition by translocation-defective RT inhibitor 4’-ethynyl-2-fluoro-2’-deoxyadenosine (EFdA). Proceedings of the National Academy of Sciences, 113(33):9274–9279, 2016.

23. Yuki Takamatsu, Debananda Das, Satoru Kohgo, Hironori Hayashi, Nicole S Delino, Stefan G Sarafianos, Hiroaki Mitsuya, and Kenji Maeda. The high genetic barrier of EFdA/MK-8591 stems from strong interactions with the active site of drug-resistant HIV-1 reverse transcriptase. Cell chemical biology, 25(10):1268–1278, 2018.

24. Merck Announces Clinical Holds on Studies Evaluating Islatravir for the Treatment and Prevention of HIV-1 Infection - Merck.com https://www.merck.com. Accessed on Fri, March 10, 2023.

25. Trials of long-acting islatravir for HIV treatment and prevention placed on hold https://www.aidsmap.com/news/dec-2021/trials-long-acting-islatravir-hiv-treatment-and-prevention-placed-hold. Accessed on Fri, March 10, 2023.

26. Merck Provides Update on Phase 2 Clinical Trial of Once-Weekly Investigational Combination of MK-8507 and Islatravir for the Treatment of People Living with HIV-1 - Merck.com https://www.merck.com. Accessed on Fri, March 10, 2023.

27. Merck restarts islatravir HIV treatment studies, but abandons monthly PrEP https://www.merck. com. Accessed on Fri, March 10, 2023.

28. Randolph P. Matthews, Dirk Schurmann, Deanne Jackson Rudd, Vanessa Levine, Sabrina Fox-Bosetti, Sandra Zhang, Martine Robberechts, Inge De Lepeleire, Andreas Huser, Daria J. Hazuda, Marian Iwamoto,, and Jay A. Single doses as low as 0.5 mg of the novel NRTTI MK-8591 suppress HIV for at least seven days. IAS 2017: Conference on HIV Pathogenesis Treatment and Prevention, July 2017, Paris, France.

29. Randolph P. Matthews, Xiaowei Zang, Stephanie E. Barrett, Athanas Koynov, Adrian Goodey, Tycho Heimbach, Vanessa L. Weissler, Carlien Leyssens, Tom Reynders, Zhiqing Xu, Sylvie Rottey, Ryan Vargo, Michael N. Robertson, S. Aubrey Stoch, and Marian Iwamoto. A Randomized Double-Blind, Placebo-Controlled, Phase 1 Trial of Radiopaque Islatravir-Eluting Subdermal Implants for Pre-exposure Prophylaxis Against HIV-1 Infection. JAIDS Journal of Acquired Immune Deficiency Syndromes, 92(4):310–316, nov 2022.

30. Deanne Jackson Rudd, Saijuan Zhang, Kerry L. Fillgrove, Sabrina Fox-Bosetti, Randolph P. Matthews, Evan Friedman, Danielle Armas, S. Aubrey Stoch, and Marian Iwamoto. Lack of a Clinically Meaningful Drug Interaction Between the HIV-1 Antiretroviral Agents Islatravir Dolutegravir, and Tenofovir Disoproxil Fumarate. Clinical Pharmacology in Drug Development, 10(12):1432–1441, oct 2021.

31. Stephanie E. Barrett, Ryan S. Teller, Seth P. Forster, Li Li, Megan A. Mackey, Daniel Skomski, Zhen Yang, Kerry L. Fillgrove, Gregory J. Doto, Sandra L. Wood, Jose Lebron, Jay A. Grobler, Rosa I. Sanchez, Zhen Liu, Bing Lu, Tao Niu, Li Sun, and Marian E. Gindy. Extended-Duration MK-8591-Eluting Implant as a Candidate for HIV Treatment and Prevention. Antimicrobial Agents and Chemotherapy, 62(10), oct 2018.

32. Sulav Duwal, Christof Schütte, and Max von Kleist. Pharmacokinetics and Pharmacodynamics of the Reverse Transcriptase Inhibitor Tenofovir and Prophylactic Efficacy against HIV-1 Infection. PLoS ONE, 7(7):e40382, jul 2012.

33. Sulav Duwal and Max von Kleist. Top-down and bottom-up modeling in system pharmacology to understand clinical efficacy: An example with NRTIs of HIV-1. European Journal of Pharmaceutical Sciences, 94:72–83, oct 2016.

34. S Duwal, V Sunkara, and M von Kleist. Multiscale Systems-Pharmacology Pipeline to Assess the Prophylactic Efficacy of NRTIs Against HIV-1. CPT: Pharmacometrics & Systems Pharmacology, 5(7):377–387, jul 2016.

35. Lanxin Zhang, Junyu Wang, and Max von Kleist. Numerical approaches for the rapid analysis of prophylactic efficacy against HIV with arbitrary drug-dosing schemes. PLOS Computational Biology, 17(12):e1009295, dec 2021.

36. Matthews; Randolph P, Rudd; Deanne Jackson, Zhang; Saijuan, Fillgrove; Kerry L, Sterling; Laura M, Grobler; Jay A, Vargo; Ryan C, Stoch; S Aubrey, and Iwamoto; Marian. Safety and pharmacokinetics of once-daily multiple-dose administration of islatravir in adults without hiv. JAIDS Journal of Acquired Immune Deficiency Syndromes, 88(3):314–321, 2021.

37. Jay A. Grobler, Ming-Tain Lai, Stephanie E. Barrett, Marian Gindy, Kerry Fillgrove, Wendy Ankrom, Sandra Wood, Evan Friedman, and Marian Iwamoto. Islatravir PK threshold & dose selection for monthly oral HIV-1 PrEP. Conference on Retroviruses and Opportunistic Infections (CROI), March 2021, Virtual Meeting.

38. Kathleen E. Squires, Todd A. Correll, Michael N. Robertson, Stephanie O. Klopfer, Peggy May Tan Hwang, Yun-Ping Zhou, Elizabeth Martin, and Elizabeth G. Rhee. Effect of Islatravir on total lymphocyte and lymphocyte subset count. Conference on Retroviruses and Opportunistic Infections (CROI), February 2023, Seattle, Washington.

39. Jay A. Grobler, Ming-Tain Lai, Stephanie E. Barrett, Marian Gindy, Kerry Fillgrove, Wendy Ankrom, Sandra Wood, Evan Friedman, and Marian Iwamoto. Modeling and Simulation to Optimize Islatravir Once Daily (QD) Doses in HIV Treatment Naive and Virologically Suppressed Populations. HIV Glasgow 2022, October 2022, Virtual Meeting.

40. Guillemette Antoni, Cécile Tremblay, Constance Delaugerre, Isabelle Charreau, Eric Cua, Daniela Rojas Castro, François Raffi, Julie Chas, Thomas Huleux, Bruno Spire, et al. On-demand pre-exposure prophylaxis with tenofovir disoproxil fumarate plus emtricitabine among men who have sex with men with less frequent sexual intercourse: a post-hoc analysis of the anrs ipergay trial. The lancet HIV, 7(2):e113–e120, 2020.

41. Jose A. Lebron; Zhanna Sobol; John Barnum; Tracy L. Diamond; Daphne Ma; Bhavana Bhatt; Nianyu Li; Zhibin Wang; Bang-Lin Wan; Fangbiao Li; Midhun Korrapati; Yuan Li1; Qian Huang; Kerry L. Fillgrove1; Qiuwei Xu; Amy G. Aslamkhan; Liping Liu; Morgan Monslow; Ryan Staupe; Jia Yao Phuah; Nithya Thambi; Kalpit A. Vora1; Ryan Vargo; Ernest Asante-Appiah and Sandrine Ferry-Martin. Investigational studies to understand the decreases in lymphocytes seen clinically with Islatravir (ISL) and enabling 1the initiation of new clinical trials. 19th European AIDS Conference, October 2023, Warsaw, Poland.

42. Kakuda; Thomas N. Pharmacology of nucleoside and nucleotide reverse transcriptase inhibitor-induced mitochondrial toxicity. Clinical therapeutics, 22(6):685–708, 2000.

43. Holec; Ashley D, Mandal; Subhra, Prathipati; Pavan K, and Destache; Christopher J. Nucleotide reverse transcriptase inhibitors: a thorough review, present status and future perspective as hiv therapeutics. Current HIV research, 15(6):411–421, 2017.

44. Peter L Anderson, Thomas N Kakuda, and Kenneth A Lichtenstein. The cellular pharmacology of nucleoside-and nucleotide-analogue reverse-transcriptase inhibitors and its relationship to clinical toxicities. Clinical Infectious Diseases, 38(5):743–753, 2004.

45. Adrian S Ray. Intracellular interactions between nucleos (t) ide inhibitors of HIV reverse transcriptase. AIDS Rev, 7(2):113–125, 2005.

46. Susan-Resiga; Delia, Bentley; Alice T, Lynx; Matthew D, LaClair; Darcy D, and McKee; Edward E. Zidovudine inhibits thymidine phosphorylation in the isolated perfused rat heart. Antimicrobial agents and chemotherapy, 51(4):1142–1149, 2007.

47. Caitlin Shepard, Joella Xu, Jessica Holler, Dong-Hyun Kim, Louis M Mansky, Raymond F Schinazi, and Baek Kim. Effect of induced dNTP pool imbalance on HIV-1 reverse transcription in macrophages. Retrovirology, 16(1):1–11, 2019.

48. Anderson; Peter L, Marzinke; Mark A, and Glidden; David V. Updating the adherence–response for oral emtricitabine/tenofovir disoproxil fumarate for human immunodeficiency virus pre-exposure prophylaxis among cisgender women. Clinical Infectious Diseases, 76(10):1850–1853, 2023.

49. Mia Moore, Sarah Stansfield, Deborah J Donnell, Marie-Claude Boily, Kate M Mitchell, Peter L Anderson, Sinead Delany-Moretlwe, Linda-Gail Bekker, Nyaradzo M Mgodi, Connie L Celum, et al. Efficacy estimates of oral pre-exposure prophylaxis for hiv prevention in cisgender women with partial adherence. Nature medicine, pages 1–5, 2023.

50. Randolph P. Matthews, Deanne J. Rudd, Vanessa Levine, Sandra Zhang, Laura Sterling, Jay Grobler, Ryan Vargo, S. Aubrey Stoch, and Marian Iwamoto. Multiple daily doses of MK-8591 as low as 0.25 mg are expected to suppress HIV. Conference on Retroviruses and Opportunistic Infections (CROI), March, 2018, Boston, Massachusetts.

51. Craig W. Hendrix, Sharon Hillier, Linda-Gail Bekker, Sharlaa Badal-Faesen, Sharon A. Riddler, Edward J. Fuchs, Pippa Macdonald, Stacey Edick, Rebeca M. Plank, Cynthia M. Chavez-Eng, Barbara Evans, Xiaowei Zang, Ryan C. Vargo, Michael N. Robertson, and Munjal Patel. Islatravir distribution in mucosal tissues, PBMC & plasma after monthly oral dosing. Conference on Retroviruses and Opportunistic Infections (CROI), Abstract 83, February 2022, Virtual Conference.

52. Jessica E Haberer, David R Bangsberg, Jared M Baeten, Kathryn Curran, Florence Koechlin, K Rivet Amico, Peter Anderson, Nelly Mugo, Francois Venter, Pedro Goicochea, et al. Defining success with hiv pre-exposure prophylaxis: a prevention-effective adherence paradigm. *AIDS (London*, England), 29(11):1277, 2015.

53. Sulav Duwal, Laura Dickinson, Aye Khoo, and Max von Kleist. Mechanistic framework predicts drug-class specific utility of antiretrovirals for HIV prophylaxis. PLoS Comput Biol, 15(1):e1006740, jan 2019.

